# A New Method to Detect Event-Related Potentials Based on Pearson’s Correlation

**DOI:** 10.1101/022046

**Authors:** William Giroldini, Luciano Pederzoli, Marco Bilucaglia, Simone Melloni, Patrizio Tressoldi

## Abstract

Event-Related Potentials (ERPs) are widely used in Brain-Computer Interface applications and in neuroscience.

Normal EEG activity is rich in background noise and therefore, in order to detect ERPs, it is usually necessary to take the average from multiple trials to reduce the effects of this noise.

The noise produced by EEG activity itself is not correlated with the ERP waveform and so, by calculating the average, the noise is decreased by a factor inversely proportional to the square root of N, where N is the number of averaged epochs.

This is the easiest strategy currently used to detect ERPs, which is based on calculating the average of each ERP’s waveform, these waveforms being time-and phase-locked.

In this paper a new method called GW6 is proposed, which calculates the ERP using a mathematical method based only on Pearson's Correlation.

This results in a graph with the same time resolution as the classical ERP and which contains only positive peaks representing the increase -in consonance to the stimuli - in EEG signal correlation over all channels.

This new method is also useful for selectively identifying and highlighting any hidden components of the ERP response that are not phase-locked, and that are usually hidden in the standard and simple method based on the averaging of all the epochs.

These hidden components seem to be caused by variations (between each successive stimulus) of the ERP's inherent phase latency period (jitter), although the same stimulus across all EEG channels produces a reasonably constant phase.

For this reason, this new method could be very helpful to investigate these hidden components of the ERP response and to develop applications for scientific and medical purposes.

Moreover, this new method is more resistant to EEG artifacts than the standard calculations of the average.

The method we are proposing can be directly used in the form of a process written in the well known Matlab programming language and can be easily and quickly written in any other software language.

## Introduction

The Event-Related Potential (ERP) is an electroencephalographic (EEG) signal recorded from multiple brain areas, in response to a single short visual or auditory stimulus or muscle movement [12, 18, 2, 10].

ERPs are widely used in Brain Computer Interface (BCI) applications and in neurology and psychology for the study of cognitive processes, mental disorders, attention deficit, schizophrenia, autism, etc.

ERPs are weak signals compared to spontaneous EEG activity, with very low signal-to-noise ratio (SNR) [12], and are typically comprised of two to four waves of low amplitude (4–10 microvolts) with a characteristic positive wave called P300, which has a latency period of about 300 milliseconds in response to the stimulus. The detection of ERPs is an important problem and several methods exist to distinguish these weak signals. Indeed, ERP analysis has become a major part of brain research today, especially in the design and development of BCIs [19].

In this paper, the definition and description of the ERP is focused mainly on the P300 because it is the simplest way to present our new ERP detection method.

We will not be considering fast evoked potentials (EVP), such as the brainstem auditory EVP, because they require a fast sampling rate (around 1000 Hz), an averaging of perhaps 1000 responses and an upper frequency filtering of about 100 to 1000 Hz.

Since the ERP is considered a reproducible response to a stimulus, with relatively stable amplitude, waveform and latency, the standard method to extract ERPs is based on the repeated presentation of the stimulus about 90–100 times, with a random inter-stimulus time of a few seconds. This strategy allows calculating the ERPs by averaging several epochs that are time-locked and phase-locked.

The averaging method is based on the assumption that the noisy EEG activity is uncorrelated with the ERP waveform, and consequently calculating the average decreases the noise by a factor of 1/SQR(N) (inverse of square root of N), where N is the number of averaged epochs. Since the background EEG activity has a higher amplitude than the ERP waveform, the technique of averaging highlights the ERPs and reduces the noise. This is the easiest strategy currently used to detect ERPs, and is also used in this paper as a reference method to be compared with our new method (called GW6) to calculate ERPs.

In general, to calculate ERPs by the method of averaging, essentially three conditions or hypotheses must be satisfied:

1. The signal is time-locked and waveform-locked.
2. The noise is uncorrelated with the signal.
3. The latency is relatively stable (low jitter).

The GW6 method also requires these three conditions, but it is less restrictive about the stability of the phase and latency, and it is also less sensitive to residual artifacts present in the EEG signals. The averaging of epochs is nevertheless only the last step in the calculation of the ERP.

Several pre-processing stages are usually necessary because the EEG signals are prone effects from important artifacts such as eye movements, heartbeat (ECG artifacts), head movements, bad electrode-skin contacts, etc. All these artifacts can be several times larger (up to 10–20 times or more) than the underlying ERPs, therefore they are able to alter calculated averages with random waves and peaks which can hide the true ERP waveform.

The first most used pre-processing step includes a band-pass filter in the range of 1 to 20 Hz obtained with a digital filter, which must not change the signal phase. The Reverse Fourier Transform is suitable for this purpose, among other methods. Many researchers have suggested that the P300 component is primarily formed by transient oscillatory events in the range which includes delta, theta and alpha, and therefore a 1 to 14 Hz band-pass could be sufficient [22].

The successive step includes a variety of methods: among the most used is the Independent Component Analysis (ICA) algorithm [15, 20], which allows separating the true EEG signal from its undesirable components (twitches, heartbeat, etc.). In general, this method requires a decision on which signal component (after separation) is to be chosen.

One of the most common problems is the removal of ocular artifacts from the EEG signals, for which purpose several techniques were developed based on the subtraction of the averaged electrooculograms and also on autoregressive modeling or adaptive methods [9, 6].

Blind Source Separation [11] is a technique based on the hypothesis that the observed signals from a multichannel recording are generated by a mixture of several distinct source signals. Using this method, it is possible to isolate the original source signal by applying some kind of transformation to the set of observed signals.

Discrete Wavelet Transform is another method that can be used to analyze the temporal and spectral properties of non-stationary signals [21, 16, 8].

The artificial neural network, known as Adaptive Neuro-Fuzzy Inference System, was described as useful for P300 detection [17]. Moreover, the Adaptive Noise Canceller can also detect ERPs [1].

So-called “winsorization” is a method which reduces the effects of blinking, voluntary and involuntary eye movements, muscle activity, or the subject’s movements, all of which can cause anomalies in the recorded signal. To reduce the effects of such artifacts, the data from each channel are processed and the amplitude values exceeding a lower and a higher percentile threshold are replaced respectively by the lower and upper percentile [7].

A good description of the ERP technique and waves components is made by Steven J. Luck [14].

## Materials and Methods

In this paper, the GW6 method is described step-by-step, as well as using a procedure written in Matlab programming language (see APPENDIX). Ultimately, this method is applied to true EEG signals recorded using a low-cost EEG device, the Emotiv EPOC® EEG Neuroheadset. This is a wireless headset and consists of 14 active electrodes and 2 reference electrodes, located and labeled according to the international 10–20 system. Channel names are: AF3, F7, F3, FC5, T7, P7, O1, O2, P8, T8, FC6, F4, F8 and AF4. The acquired EEG signals are transmitted wirelessly to the computer by means of weak radio signals in the 2.4 GHz band. The Emotiv’s sampling frequency is 128 Hz for every channel and the signals are encoded with a 14-bit definition.

Moreover, the Emotiv hardware operates preliminarily on signals with a band-pass filter from 0.1 to 43 Hz, consequently the output signals are relatively free from disturbances caused by the 50 Hz power-line frequency, however they are often rich in artifacts.

The Emotiv EPOC® headset was successfully used to record ERPs (Badcock et al., 2013) although it is not considered a medical-grade device. Emotiv EPOC® was moreover widely used for several researches in the field of Brain Computer Interface (BCI) [5, 13].

We collected and recorded the raw signals from the Emotiv EPOC® headset using software we created ourselves and a special data-type format based on the.CSV format. The same software we created was used to give the necessary auditory and/or visual stimuli to the subject.

Participants were ten healthy volunteers, ranging in age from 28 to 69 years, informed in advance about the experimentation’s purpose. Each participant gave written consent, with IRB (Institutional-Review-Board) approval. Participants had normal vision and no history of hearing-related problems; they were resting in a comfortable position during the tests.

The ERPs were induced by an auditory stimulus (pure 500 Hz sine wave) with a simultaneous light flash using an array of 16 red high-efficiency LEDs. The stimulus length was 1 second, and the stimuli were repeated from 100 up to 128 times.

Using the original EEG (mastoidal) reference electrode of the Emotiv EPOC® headset, we recorded a first group of experimental data on 14 channels. Another group of better quality EEG files were recorded with the reference electrodes connected to the earlobes, a variation that assures better quality of the signals, rather than in the standard configuration of the Emotiv EPOC® headset where the reference electrodes are placed on an active area of the head.

## The new algorithm

With our software we pre-processed the EEG files using digital data-filtering in the 1 to 20 Hz band followed by a method we called Normalization.

The filtering was performed using the Reverse Fourier Transform which does not change the signal phase. The conservation of the original phase of signals is very important for the application of our method. On the other hand, the conservation of the information about the phase pattern of the signals, rather than the simple power of the signals, was found important also in the representation of semantic categories of objects, especially in the low-frequency band (1 to 4 Hz) [4].

The second step in pre-processing was signal Normalization: the signals from each channel, i.e.

S(x), comprised of 4 second epochs, were normalized as:

S(x) = K_*_[S(x)-mean]/Std

where S(x) is firstly reduced to a zero-mean signal, where mean is the mean value of the signal in the epoch; Std is the corresponding standard deviation and K is an experimental constant which restores the averaged optimized amplitude of the EEG signal. The K factor is the Standard Deviation of a good quality EEG signal, found experimentally using these specific instruments. This number was calculated as K = 20. This normalization step created an epoch with a shape identical to that of the original EEG signal, but transferred into a uniform scale, with comparable amplitude for every epoch. The entire file fully processed as such was saved as new file in.CSV format, containing all the information about the start and the end of each stimulus.

Note that it is also possible to pre-process only time-locked epochs, for example 3 seconds long, corresponding to each stimulus [pre-stimulus + stimulus (1 s) + post-stimulus], and in general this procedure gives non-identical results although very similar to the previous method based on the filtering and saving of the entire file.

Another common way to pre-process the data for the ERP calculation is the exclusion of every epoch with an amplitude above a fixed threshold, for example 80 microvolts. A disadvantage of this technique is that a large number of epochs could be discarded and consequently the average could be calculated on insufficient data. In our software, we also used this procedure to eliminate epochs with strong artifacts above 100 microvolts still present after the digital filtration.

In this paper, we will to illustrate a new method which is useful for detecting ERPs even among particularly noisy signals and with significant latency variations, known as “latency jitter”.

Our method, called GW6, is less restrictive regarding the issue of jitter. It also allows ERP detection when the standard approach, based on the average, fails or gives unsatisfactory results due to several artifacts. However the GW6 method does not reproduce the typical biphasic waveform of the ERP but rather an always positive waveform. For this reason, this new procedure is useful if used together with the classic technique of averaging, rather as an alternative to the latter.

The GW6 method uses Pearson's Correlation extensively for all EEG signals recorded by a multichannel EEG device. By using the method of averaging, it is possible to work with a single EEG channel too, whereas the GW 6 method works only with a multichannel EEG device, starting from a minimum of 6 channels. Nevertheless it is also possible to progressively calculate the ERP in each channel.

In many papers describing a mathematical method to analyze something, formulas are usually given, which must be subsequently translated into a computer-language, for example C, C++, Visual Basic, Java, Python, Matlab, or other. This step could be difficult and limit the release and application of some useful methods. In this paper, we will describe this new algorithm as a step-by-step procedure and also in the simple and well-known Matlab programming language, in order to ease its application (see Appendix).

We describe the basic idea of this new method in Figures 1 and 2:

**Fig. 1:**
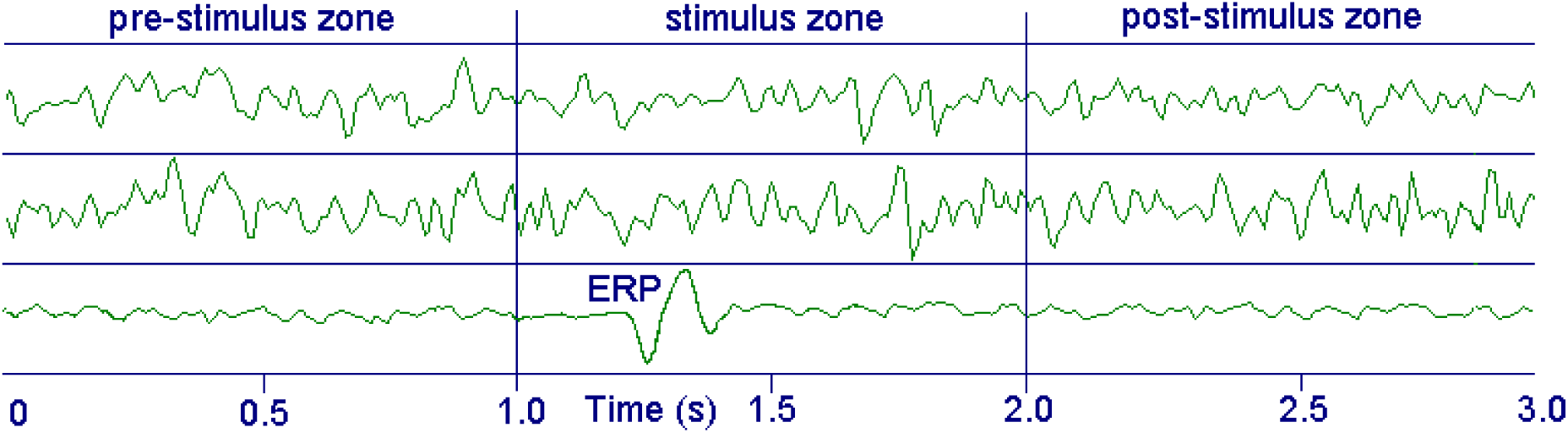
The two upper tracks represent the raw signals of two EEG channels in time-locked epochs, whereas the lower track is the average of a sufficient number (about 100) of epochs for each channel (ERP is not to scale). The figure shows a positive peak about 300 ms after the stimulus’s onset (P300 wave). The ERP’s typical duration is about 300–500 ms, depending on the kind of stimulus and band-pass filtering of the signal.

**Fig. 2:**
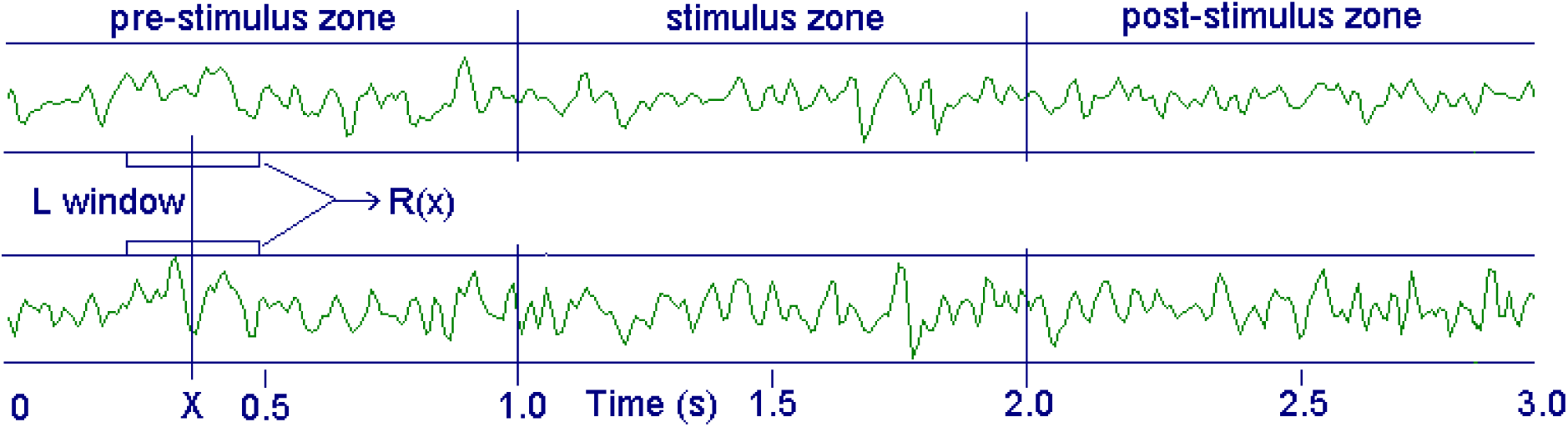
The double data-window lasting L is shifted progressively along the two tracks S1(x) and S2(x), and the corresponding Pearson's Correlation between the two windows is calculated and stored in the vector R(x), where x is the sampling data index of the tracks.

Let us now consider the Figure 2 and the double data-window lasting L (about 270 ms, 34 samples) centered at point X of the signal. We can calculate the Linear Pearson's Correlation between these two data-segments and the result will be a number *r* represented by the vector R(x), which can be calculated for every point X simply by progressively moving the windows along by one sample unit at a time (sliding windows). In general, the averaged value of R(x) will vary from the pre-stimulus zone to the stimulus zone because the (auditory or/and visual) stimulus changes the correlation between the two EEG signals, which represent the activity of different parts of the brain. An interval about 270 ms long was selected because it represents the typical amount of time required for a conscious response corresponding to the P300 wave, but different intervals could be selected for fast Evoked Potentials, or other types of stimulus.

This change of correlation can appear either as an increase or a decrease with respect to the baseline (i.e. the zone preceding the stimulus). Let us consider a real example, based on the Emotiv EPOC®, where the number of channels is NC = 14, the sampling frequency is 128 samples/second, the stimulus length is one second, and the epoch length is 3 seconds. In this case it is possible to calculate the vector R(x) in a number of pair combinations Nt = NC_*_(NC - 1)/2 = 91.

The result could be expressed using a new array R(I, X) where I = 1... 91, and X = 1... 384.

This last number arises from a 3-seconds length epoch and 128 samples/second, with the stimulus given at sample number 128, and stopped at sample number 256, after an extra second.

Each value of R(I, X) comes from the Pearson's Correlation between two data-windows of duration L (i.e. 34 samples) centered on point X, and for any pair combination of the NC channels.

Moreover, the array R(I,X) is averaged along all the Ns stimuli given to the subjects.

In general, we can represent the raw signals as a time-locked array of V(C, X, J) type, where C = 1... 14 are the EEG channels, X = 1... 384 are the samples along 3 seconds, and J = 1... Ns is the number of stimuli given to the subject, usually about 100. The entire GW6 procedure is better described in the Matlab method (see Appendix).

The following are the processing stages based on the 14-channel Emotiv EPOC® device, but not limited to this specific device (the numbers here described are only examples):

**Stage 1:** filtration of the.CSV file in the frequency band 1 - 20 Hz, Normalization and new saving of the entire file. It is however possible to omit this stage and go directly to filtration and Normalization on the time-locked epoch of each stimulus of the file.
**Stage 2:** from the raw EEG data, or from the pre-processed file, calculate all the time-locked epochs and storage of the data in the array V(C, X, J), where C = 1... 14 are the channels, X = 1… 384 are the samples, and J = 1... 100 is the stimulus index. However, due to the presence of the L windows, we need a longer array for processing, for example the length could be increased by two tails of length L = 34, giving a total number of L + 384 + L = 452 samples, with the stimulus starting at X=162 and stopping at X=290. Now the array V(C, X, J), filtered and normalized, is renamed as the new array W(C, X, J). V(C, X, J) + filtration + normalization → W(C, X, J). It is very important that any pre-processing method modifying the correlation among the signals must be excluded.
**Stage 3:** calculation of the simple average of W(C, X, J) among all Ns epochs (number of stimuli), giving the final array Ev(C, X), which is the simple and classic ERP of each channel.

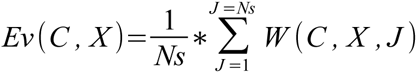 A detail to note: when processing has finished, the X index is easily recalculated in order to cut off the tail lengths L at the beginning and end, giving the final array Ev(C, X) where X = 1…384 and C = 1...14 This array Ev(C, X) is used as a comparison with the result of our method, and to show the differences in the waveform of the resulting ERP.
**Stage 4:** calculation of all the Pearson's Correlation combinations using a sliding-window 270 ms long, as described in Fig. 2. The result is the array R(I, X), with I = 1... 91, which is also calculated as the average of all the stimuli. Here too, at the end of this stage the index X is recalculated in order to cut off the initial and final L tails, giving the final array R(I, X) where I = 1. 91 and X = 1... 384 (see Appendix).
**Stage 5:** Calculation of the baseline mean value for each Nt combination (Nt = 91). The best baseline is calculated as a balanced average of pre-stimulus plus post-stimulus of each combination, then subtracting this baseline from the array R(I, X), and taking the absolute value.

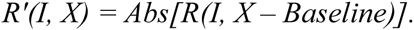
**Stage 6:** average along all the Nt combinations (and all the stimuli), giving the final array Sync1(X), which represents the global variation of the EEG correlations during a 3 second epoch.

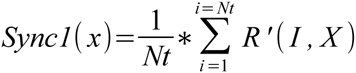

It is also possible to calculate an equivalent array Sync2(C, X) for each channel C (see Appendix). The global array Sync1(X) and Sync2(C, X) will show one or more positive peaks in the ERP zone, as shown in Fig. 3, and these peaks represent the variations of correlation among the different brain zones (electrodes) during the stimulus.

**Fig. 3:**
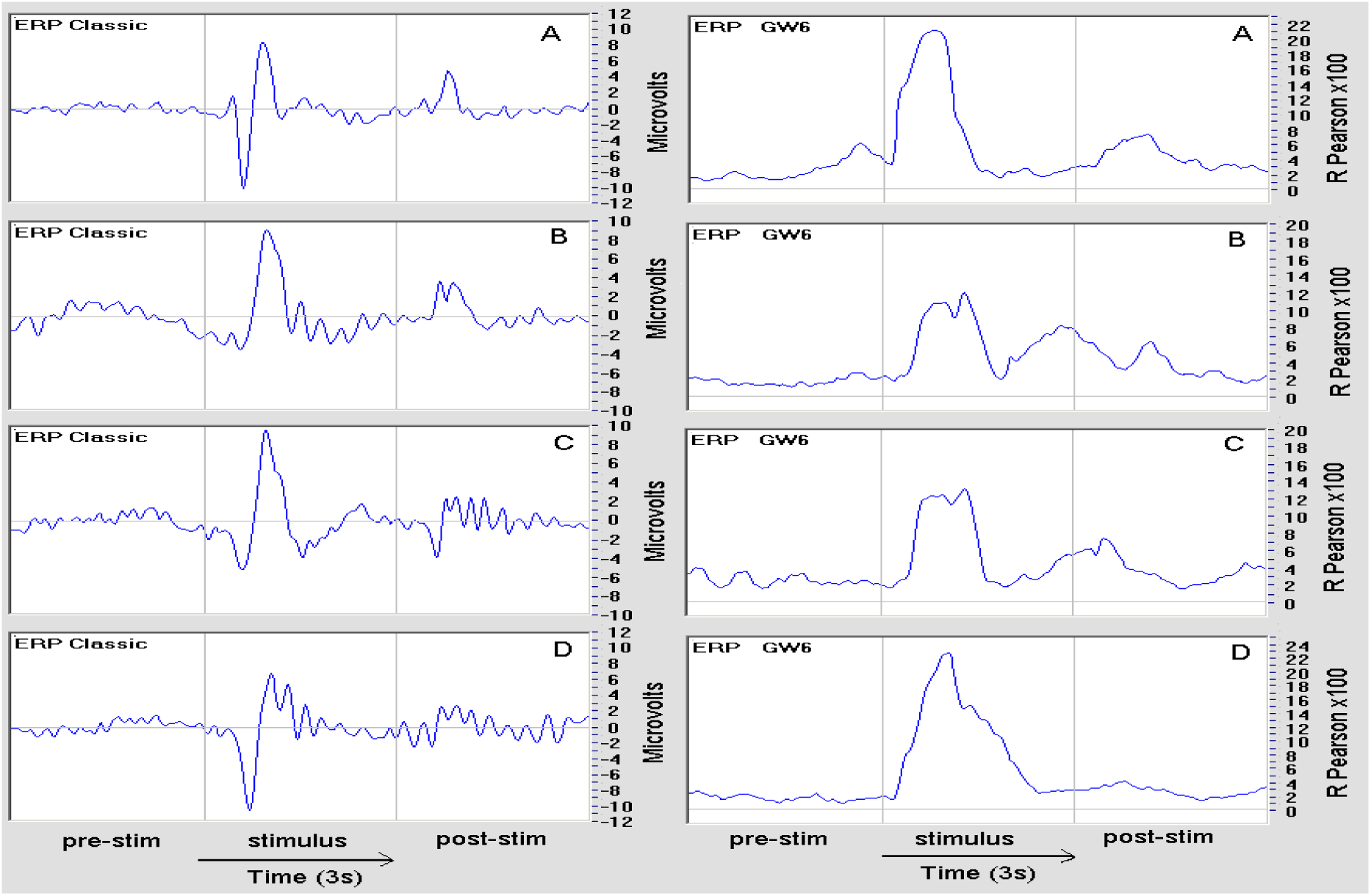
In these pictures, shown as examples, the left presents the classic ERP (amplitude in microvolts). On the right is shown the corresponding GW6 graph; the result is expressed as the R-Pearson value multiplied by 100. All these graphics are the global average of 14 EEG channels and about 120 stimuli; the EEG data were filtered in the 1–20 Hz band and submitted to the Normalization routine. In all cases a positive peak is observed coinciding with the P300 maximum peak, but in the majority of cases the positive peak of the GW6 graph is larger than the corresponding classic ERP (see, for example, cases B, C and D).

## Experimental results

These graphs are examples of the typical results provided by this method:

In order to better investigate the properties of the GW6 method, we wrote an emulation software. In this software, a simple artificial ERP waveform was added to a random noise, and suitably filtered (low-pass filter) in order to reproduce the typical frequency distribution of the EEG signal. The artificial ERP signal was mixed with a variable amount of this random signal and the result was submitted both to the classic average and to the GW6 routine (Fig. 4).

**Fig. 4:**
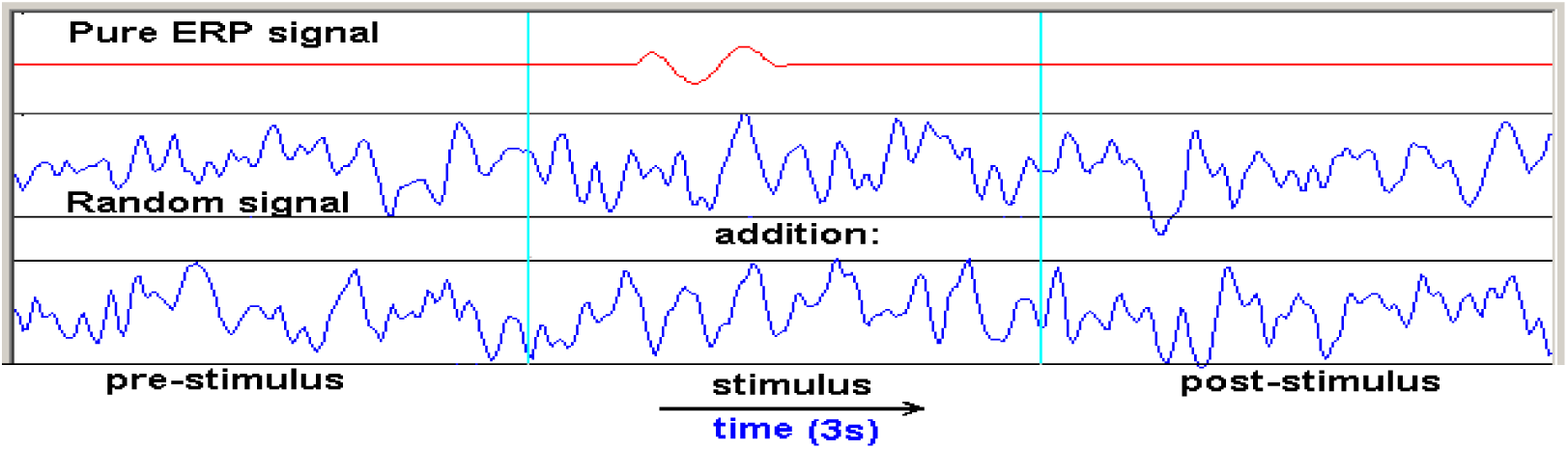
Artificial ERP signal mixed with a variable amount of a random signal and submitted both to the classic average and to the GW6 routine.

Fig. 5 shows the results of the classic average method and of the GW6 method for a progressive increase of the noise-to-signal ratio, as an average of 100 ERPs on a single channel. While the final amplitude of the ERP waveform does not change, but instead becomes progressively noisier, the GW6 graph’s amplitude (red curve) progressively drops, but with a stable residual noise.

**Fig. 5:**
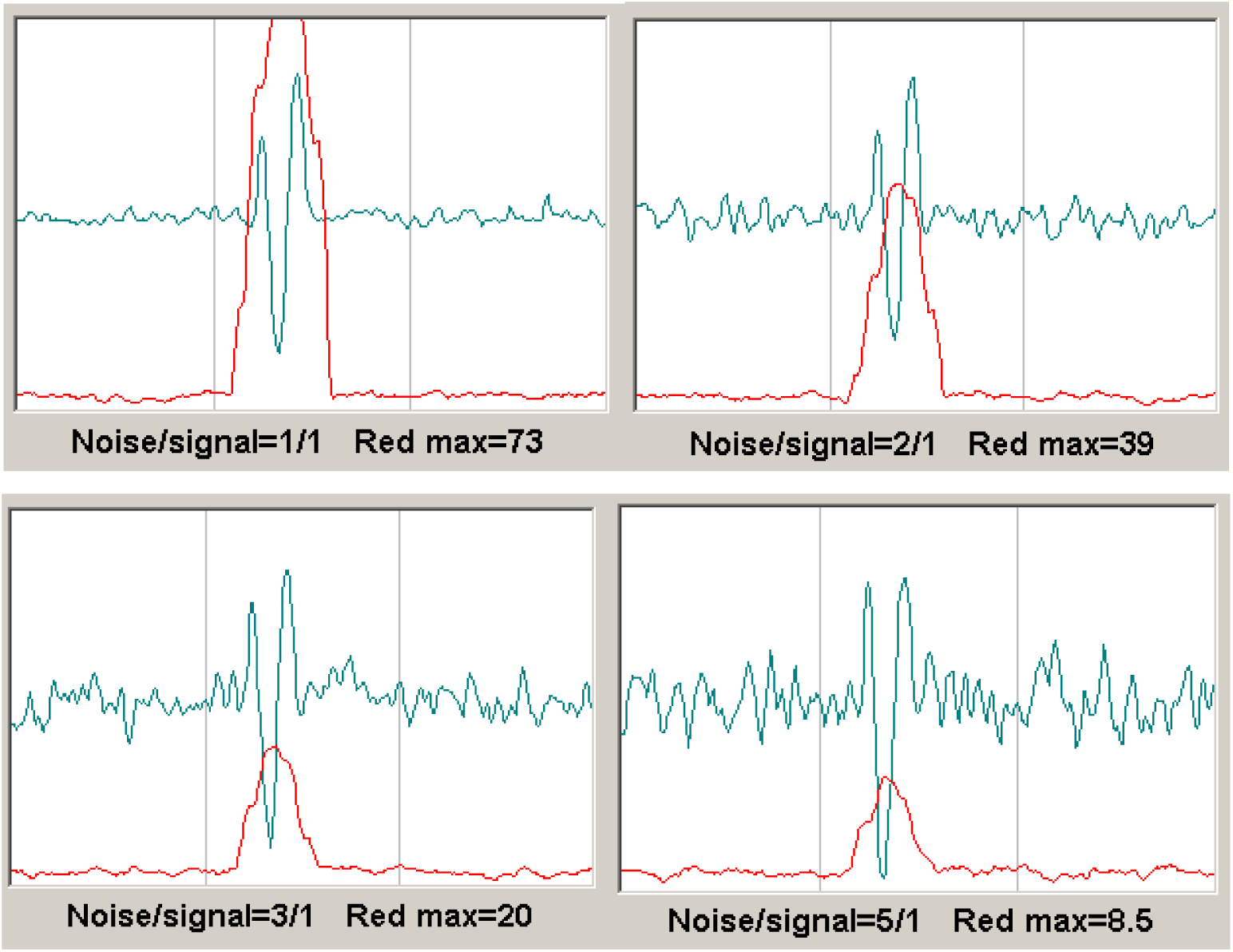
Example of the ERP + Random signal emulation for four levels of noise-to—signal ratio.

Of particular interest is the emulation of these two methods in the presence of the so-called “latency jitter”, which is an unstable ERP time latency that in some cases could affect the ERPs.

When the ERP latency is stable (Fig. 6, left picture), its average is stable too and is at its maximum amplitude. Nevertheless, if latency jitter is present (due to some physiological cause) the corresponding average decreases because each ERP does not combine with the same phase and consequently ERPs have a tendency to cancel each other out. This effect is more pronounced as the jitter increases. In the software emulation of Fig. 7 a stable noise-to-signal ratio (3/1) was used, but with a random jitter progressively increased. Moreover, the jitter was random between the ERPs, but was constant for all the channels in each ERP. The results show that the GW6 routine is more resistant to jitter than the simple classic average.

**Fig. 6:**
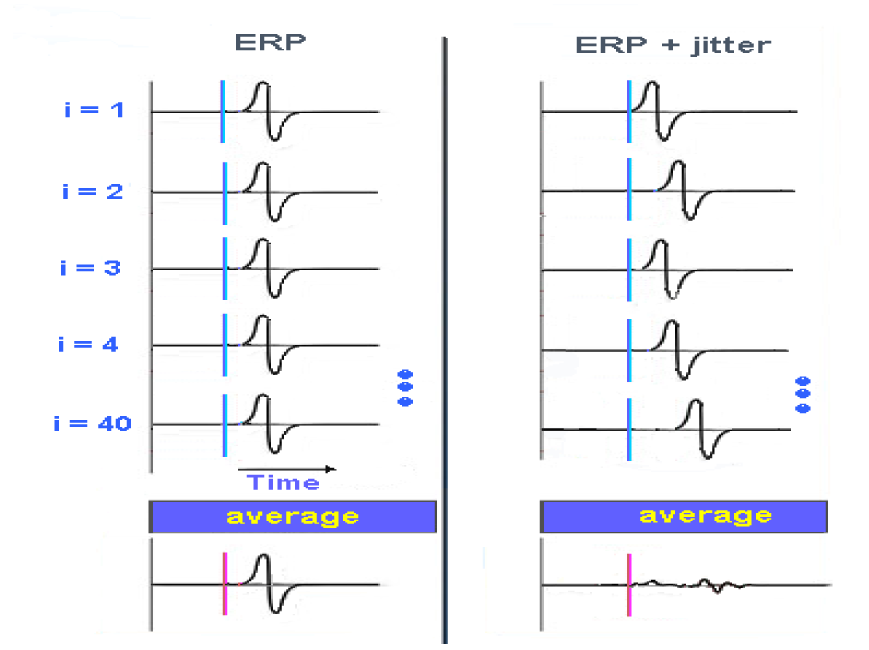
ERP with stable latency on the left and with latency jitter on the right

**Fig. 7:**
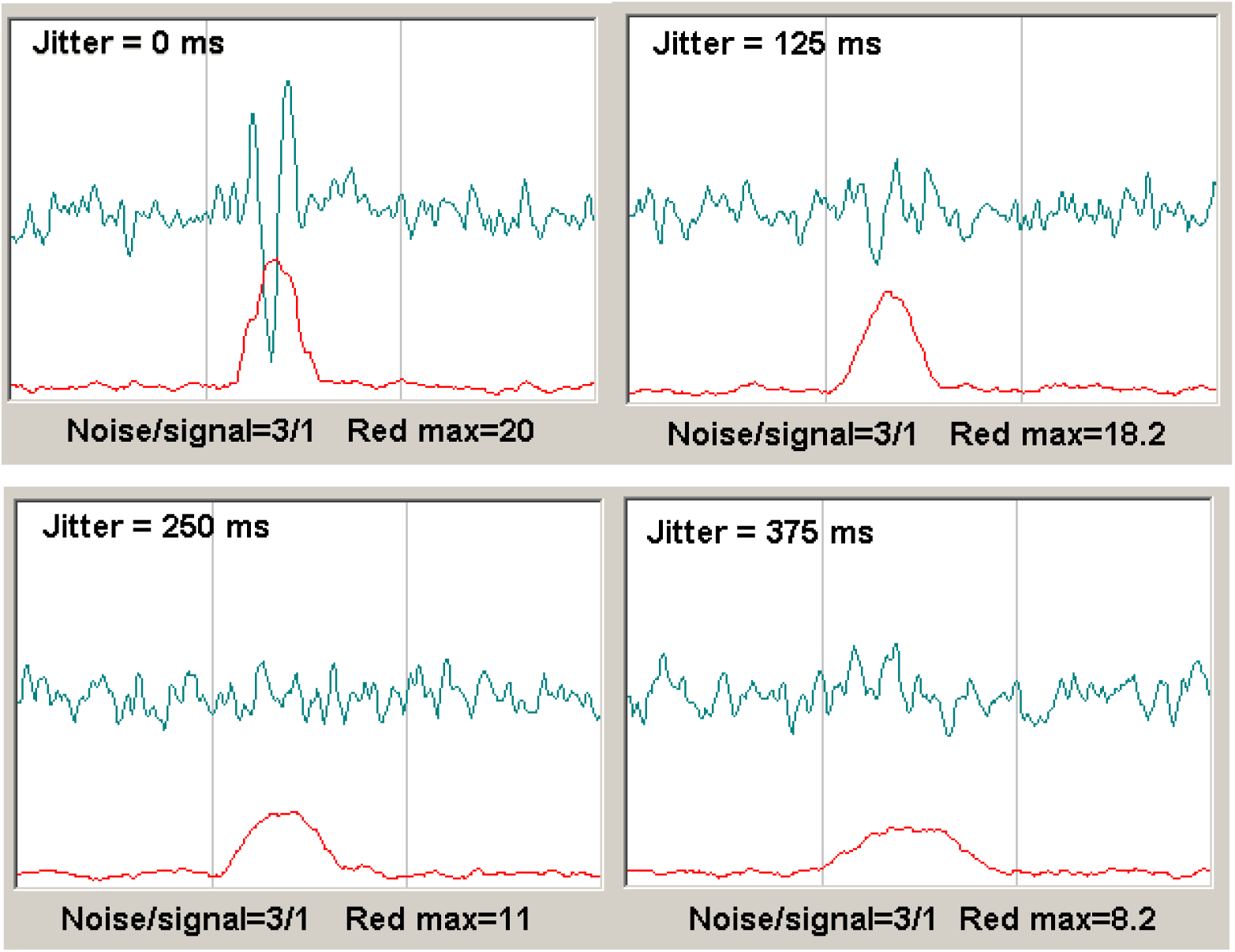
A stable noise-to-signal ratio (3/1), but with a random jitter progressively increased.

Whereas the classic ERP waveform disappears rapidly as the jitter increases, the GW6 routine gives a still identifiable result (the red curve), where the amplitude is decreases but not as rapidly, and the curve’s width is increases. This interesting property is very important, because it suggests some other possibility about the large GW6 peaks observed in Fig. 3, in particular in B, C and D cases. Following a hunch, we added a new and simple process to our software used to analyze the true ERP using both the classic and the GW6 methods. At the end of the process, which gives the typical result shown in Fig. 3, we created another procedure where the classic ERP average was subtracted (see Appendix) from the set of EEG signals W(C, X, J), giving a new array:

W'(C, X, J) = W(C, X, J) - Ev(C, X), then this new data-set was submitted to stages 3, 4, 5 and 6 previously described. Incorporating this process in our emulation software, and successively performing the same 3, 4, 5 and 6 stages, no ERP appears as a result, nor does any significant GW6 peak. This is obvious because in doing so we have canceled the ERP component from the random noise, and consequently nothing is expected to appear, but that is true only if jitter is zero (Figures 8 and 9).

**Fig. 8.**
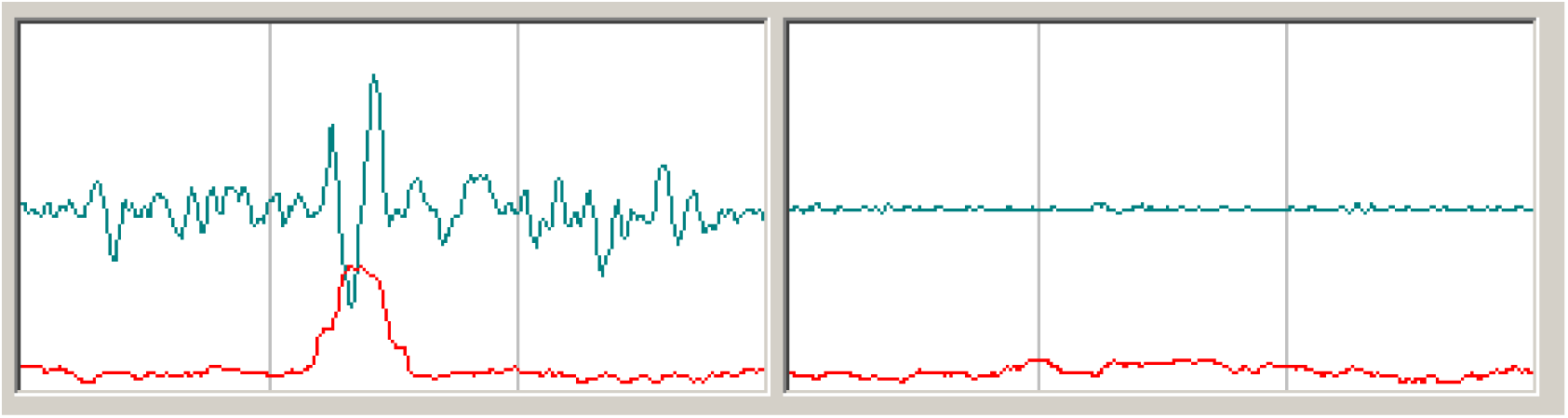
Case 1. Left: W(C, X, J) from ERP pure wave + random noise, Jitter *=* 0, average of 100 ERPs. Right: with the same processing of the corresponding W'(C, X, J) array both graphs disappear.

**Fig. 9.**
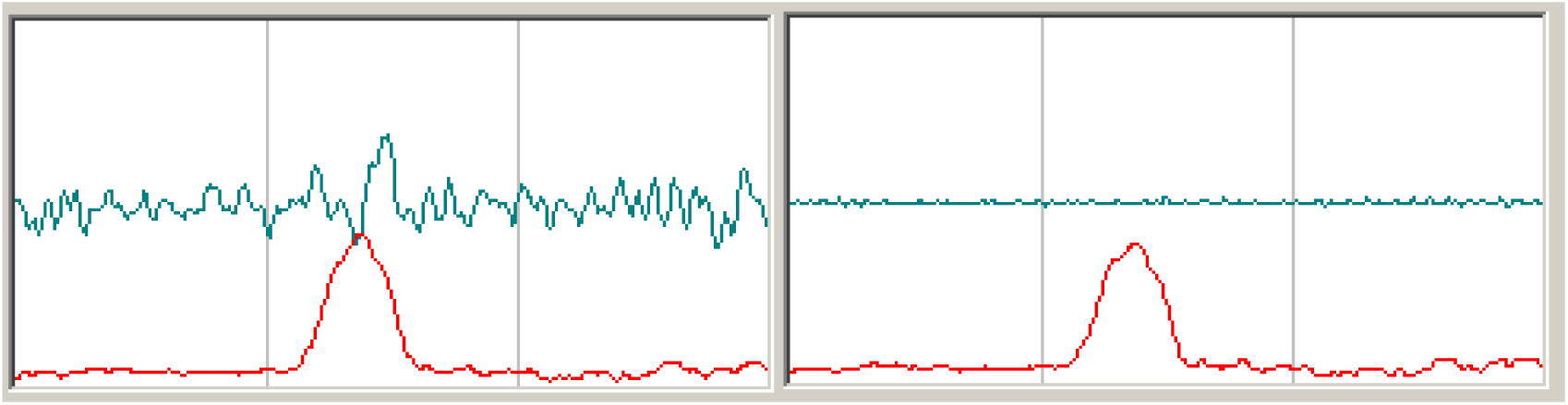
Case 2. Left: W(C, X, J) from ERP pure wave *+* random noise, Jitter *=* 78ms (from 0 to 78 ms, random), average of 100 ERPs. Right: with the same processing of the corresponding W'(C, X, J) array only the classic ERP disappears. In the presence of Jitter, the GW6 method always shows an ERP.

**Fig. 10.**
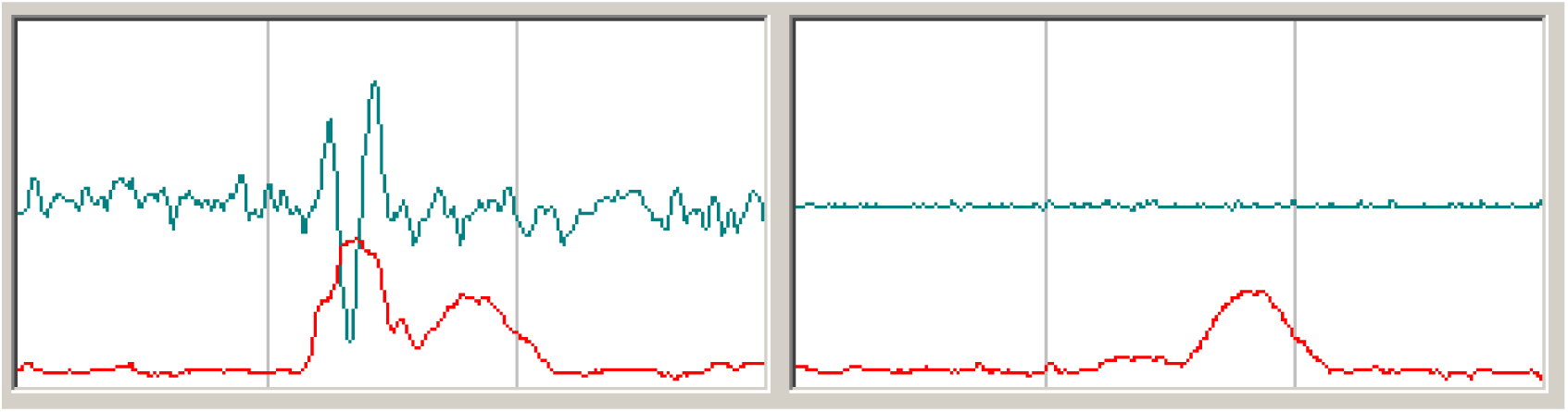
Case 3. Left: W(C, X, J) from ERP pure wave + random noise, Jitter — 0, RCS width about 400 ms, average of 100 ERPs. Right: with the same processing of the W'(C X, J) array only the classic ERP disappears, not that due to RCS.

We created a new variant in the emulation software: alongside the pure ERP wave + random signal, we also added a random common signal (RCS) to every channel only in a limited zone near the ERP, but this RCS is random between the ERPs. In this emulation variant we hypothesized that the stimulus given to the subject could not only cause a simple brain response based on a stable waveform with low jitter (the classic ERP) but also cause a non-stable waveform very similar or identical in all the EEG channels. A simple calculation of the average does not reveal this kind of electrical response, because the waveform is near random, but it is easily revealed by the GW6 method, which is based on the calculation of the variation of correlation among all the EEG channels during the stimulus.

We believe that the two last cases (4 in Fig. 11 and 5 in Fig. 12) best represent true experimental ERPs. With our emulation software many combinations and situations can be calculated. Now, if we submit our true experimental ERPs to the same procedure, i.e. analysis of the W(C, X, J) data followed by transformation into the W'(C, X, J) data-set and a new analysis, we obtain these typical results (Fig. 13):

**Fig. 11.**
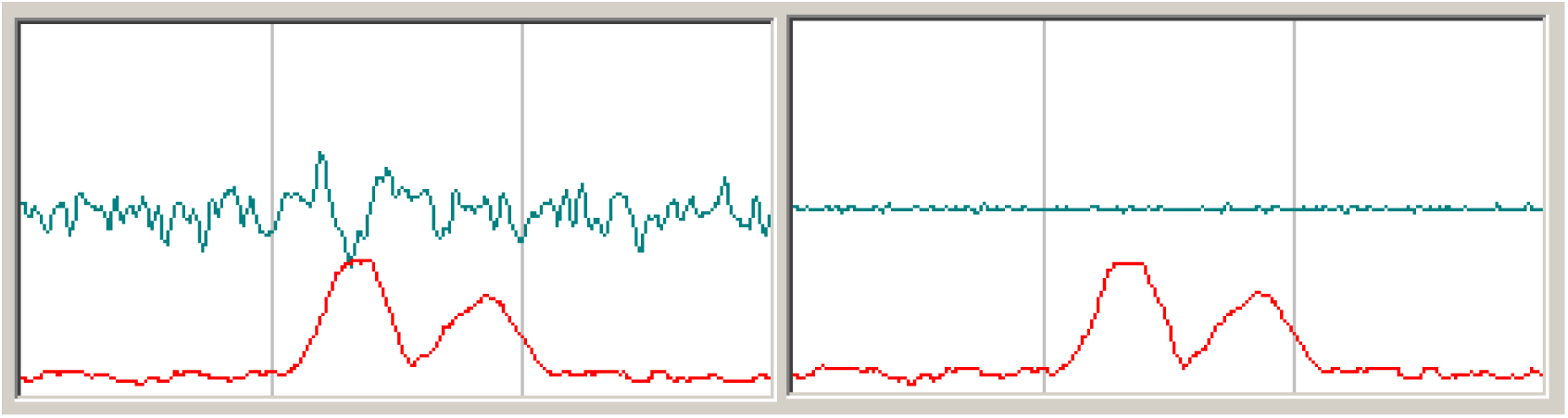
Case 4. Left: W(C, X, J) from ERP pure wave + random noise, Jitter = 78 ms, RCS width about 400 ms, average of 100 ERPs. Right: with the same processing of the W'(C, X, J) array, now both peaks are visible.

**Fig. 12.**
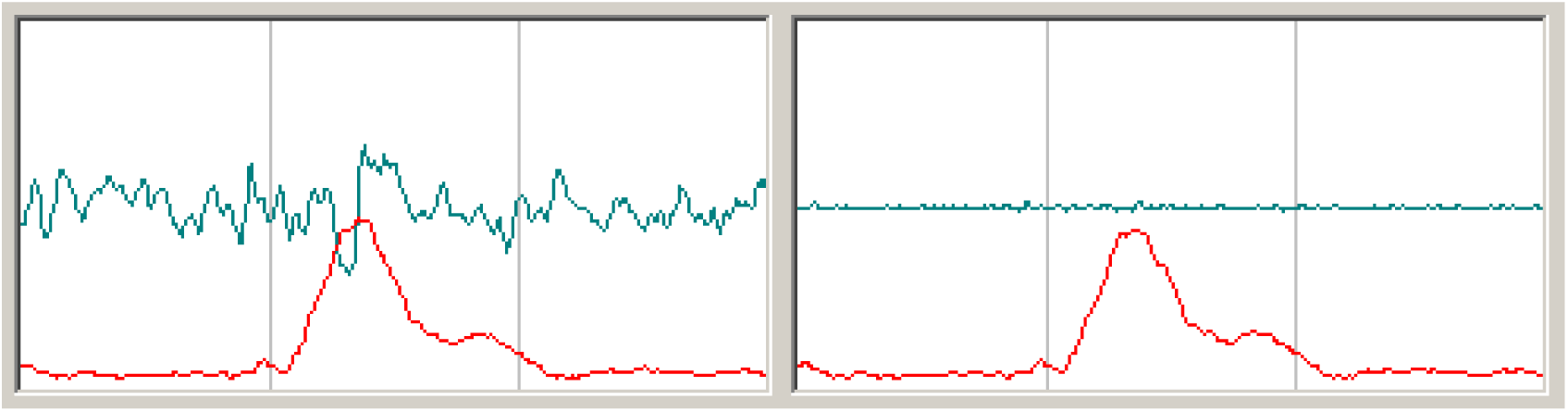
Case 5. Left: W(C, X, J) from ERP pure wave + random noise, Jitter =78ms, RCS width about 860 ms, average of 100 ERPs. Right: with the same processing of the W'(C, X, J) array, now both the peaks overlap and are visible.

**Fig. 13:**
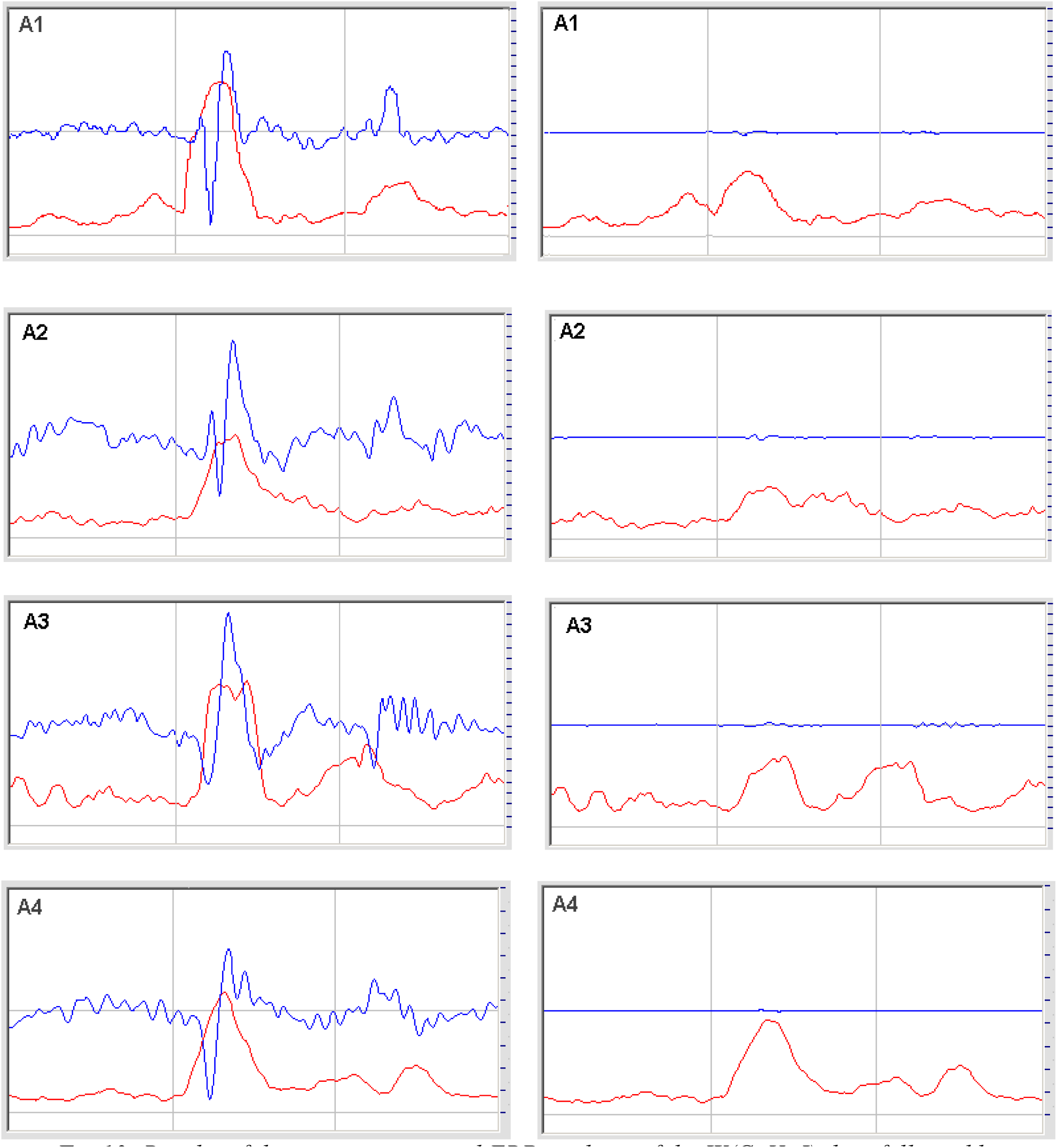
Results of the true experimental ERP analysis of the W(C, X, J) data followed by transformation into the W'(C, X, J) data-set and new method GW6.

As shown in fig. 13, in the majority of cases, after the subtraction of the classic ERP waveform from the EEG data, the GW6 method (red graph) shows a reduction in amplitude corresponding to the standard ERP wave, but other peaks are hardly changed at all, and in several cases there is minimal change to the whole graph.

## Discussion

This new method allows the calculation of ERPs as variations of the global correlations among all the EEG channels, with respect to the averaged pre-stimulus and post-stimulus zone.

Furthermore, the method shows significant peaks in the P300 zone larger than the peaks calculated from the standard procedure of averaging. In the presence of significant jitter (instability of latency) the new method is superior with respect to the classic one and shows significant peaks in this case too. Our experimental results suggest that, in the majority of cases, there is some amount of jitter coinciding with the classic ERP, and/or the significant presence of other components that are not phase-locked, such as those hypothesized in the emulation software, both under the classic ERP peak and in other areas of the graph. In particular, in each stimulus it is possible to find and identify non-phase-locked components of the ERP among all the stimuli, and which are phase-locked among all the EEG channels. These components could be easily calculated simply by subtracting the classic ERP from the EEG signal of each channel and re-analyzing the new data using the GW6 method. Consequently, it is now possible to obtain three types of graphs: first, the classic ERP based only on simple averaging, which shows both phase- and time-locked components; second, the GW6 ERP which shows all components including those that are non-phase locked; and third, the graph showing only the non-phase-locked components of the ERP. Our method is also inherently more resistant to artifacts because the Pearson's Correlation depends only on signal phase and not on amplitude, while the artifacts are mainly due to strong signal amplitude variations. Although this method is not compatible with all the pre-processing methods which change the correlation among EEG signals, it is applicable to the majority of cases, and probably also in cases not presented or discussed here due to limitations in our instrumentation and experimental setup.

## Conclusions

The purpose of this paper is not, at this time, to accurately investigate the EEG response to a specific stimulus or specific experimental protocol, but only to propose a new method for ERP detection and analysis that could become very important for future research about the nature, origin and characteristics of ERPs in light of the preliminary result presented here.

In particular, this new method could be very useful for investigating hidden components of the ERP response, with a possible important application for medical purposes and in the fields of neurophysiology and psychology.

Furthermore, we emphasize choosing the well-known Matlab language tool for mathematical processing so that the method can be easily used with and applied to independent software as well as Research.

## Acknowledgments

**T**his study was partially supported by the research grant n. 124/12 of Bial Foundation.

Thanks to Svetlana Kucherenko for the translation of our software from Visual Basic to Matlab.

## Appendix

Now suppose that the W(C, X, J) array represents all the data to be analyzed, where C are the channels (14 in the Emotiv Epoc), X the samples from X = 1 to 452, J = 1 to Ns, where Ns is the number of stimuli given to the subject (usually Ns = 100–130).

We must moreover suppose that the sampling frequency is 128Hz and we take in consideration a time-locked epoch of 3 second (pre-stimulus, stimulus, post-stimulus, 384 samples) plus two additional tails (the L windows, 34 samples) giving a total of 452 samples. These tails are necessary for the elaboration, but they will be cut off at the end, in order to give a final epoch of 3 seconds.

Moreover, we suppose that the stimulus starts at X = 162 and stops at X = 290.

Data in the W(C, X, J) array must be with zero-average (easy to implement, here omitted).

The unity of the W(C, X, J) data could be in microvolts, or raw data at 10, 12, 16 bit, etc.

% The essential core of GW6.m routine in Matlab language

V1 = zeros(1,800);
V2 = zeros(1,800);
A1 = zeros(1,800);
A2 = zeros(1,200);
F1 = 0; F2 = 0; F3 = 0; I = 0; J =0;
XM = 0; YM = 0; Rc = 0; X = 0;
Ax = 0; Bx = 0; Np = 0; Tc = 0; C =0;
Nz = 0; A = 0; U =0; X1 = 0; X2 = 0;

% definition of the value of data
N1 = 452; % 384 samples + 2 tails of 34 samples
NC = 14;
Ns =100;
Np = 34; % 34 is a L window of about 270 ms, at 128 samples/s, could be changed
Tc = Np/2; % half window L
Nt = (NC ^ 2 - NC) / 2; % in our case Nt=91 is the number of combination with 14 channels

R = zeros(Nt+1,N1+1); % put zero each element of array R(I, X)

for J = 1: Ns % for all the stimuli given
U = 0; I = 0;
for Ax = 1: (NC - 1)
for Bx = (Ax + 1): NC
I = I + 1; % counter of the progressive combinations of two channels for U = Tc: (N1 -Tc)
X1 = U - Tc + 1;
X2 = U + Tc + 1;
A = 0;
for X = X1: X2 A = A + 1;
V1(A) = W(Ax, X, J);
V2(A) = W(Bx, X, J); end
Nz = A;
correlas; % subroutine of Pearson's Correlation
A1(U) = Rc;
A = 0;
end
for X = Tc: (N1 -Tc)
R(I, X) = R(I, X) + A1(X);
end
end
end
end
% averaging along all the Ns stimuli
for I = 1: Nt
for X = 1: N1
R(I, X) = R(I, X) /Ns;
end
end
% cutting of the two L tails in order to recover the interval of 3 seconds for I = 1: Nt
U = 0;
for X = (Np + 1): (N1 - Np)
U = U + 1;
R(I, U) = R(I, X);
end
end
N1 = U; % now N1 value is 384, equivalent to 3 seconds
% now the array R(I, X) is the output of this stage of elaboration

%Correlas: %%%‘ Pearson’s correlation subroutine
F1 = 0; F2 = 0; F3 = 0; XM = 0; YM = 0;
for X = 1: Nz;
XM = XM + V1(X);
YM = YM + V2(X);
end
XM = XM / Nz;
YM = YM / Nz;
for X = 1: Nz
F1 = F1 + (V1(X) - XM) * (V2(X) - YM);
F2 = F2 + (v1(x) - XM) ^ 2;
F3 = F3 + (V2(X) - YM) ^ 2;
end
F1 = F1 / Nz; F2 = F2 / Nz; F3 = F3 / Nz;
if (F2 == 0 || F3 == 0)
Rc = 0;
return
end
Rc = 100 * F1 / sqrt(F2 * F3); % the r of Pearson is multiplied by 100
return

% successive FinalElab.m elaboration
Bs = zeros(100);
B1 = 0; B2 = 0; C = 0; J = 0; X = 0; I = 0;
% Nt = 91 is defined in the previous elaboration
B1= 128; B2 = B1 + 128; N1 = 384;
% the stimulus zone is between X = B1 and X = B2
% for the calculation of a balanced baseline, we take the pre-stimulus zone
% (from X = 1 to B1), the second is the post-stimulus zone (from X = B2 to N1)

Sync1=zeros(N1);
Sync2=zeros(NC, N1);
A=0;
for I = 1: Nt % baseline calculation
A = 0; Bs(I) = 0;
for X = 1: B1
A = A + 1;
Bs(I) = Bs(I) + R(I, X);
end
for X = B2: N1
A = A + 1;
Bs(I) = Bs(I) + R(I, X);
end
Bs(I) = Bs(I) / A; % baseline for each combination
end
for I = 1: Nt
for X = 1: N1;
Sync1(X) = Sync1(X) + abs(R(I, X) - Bs(I));
end
end
for X = 1: N1
Sync1(X) = Sync1(X) / Nt;
end
% Now the array Sync1(X) is the average (global average) of Correlation for all the Nt combinations
% and for all the Ns number of stimuli.

% Calculation of the array Sync2(C, X) for each EEG channel
I = 0; U = 0;
for Ax = 1: (NC - 1)
for Bx = (Ax + 1): NC
I = I + 1; % counter of all the combinations of the channels
for U = 1: NC
if (U == Ax) || (U == Bx)
for X = 1: N1
Sync2(U, X) = Sync2(U, X) + abs(R(I, X) - Bs(I));
end
end
end
end
end

for C = 1: NC
for X = 1: N1
Sync2(C, X) = Sync2(C, X) / (NC-1);
end
end
% Now the array Sync2(C, X) is the Correlation for each channel.
% Each channel is the average of (NC-1) data.

% Simple example of the ClassicERP.m calculation
% NC are the EEG channels, N1 the number of samples time-locked
% if N1 = 452 as in the W(C, X, J) array, a final cutting of two tails of length L = 34 should be done
% like in the GW6.m elaboration.

Ev= zeros(NC, N1); % Ev(C, X) is the array of classic ERP for each EEG channel
for J = 1: Ns % for all the stimuli
for C = 1: NC %for all the channels
for X = 1: N1
Ev(C, X) = Ev(C, X) + W(C, X, J);
end
end
end
for C = 1: NC
for X = 1: N1
Ev(C, X) = Ev(C, X) / Ns;
end
end

% Now the array Ev(C, X) is the classic ERP.

## References

1. Ahirwal M. K., Kumar A., & Singh, G. K. (2014). Adaptive filtering of EEG/ERP through noise cancellers using an improved PSO algorithm. Swarm and Evolutionary Computation, 14, 76–91.

2. Aydin S. (2008). Comparison of basic linear filters in extracting auditory evoked potentials. Turkish Journal of Electrical Engineering, 16(2), 111–123.

3. Badcock N. A., Mousikou P., Mahajan Y., de Lissa P., Thie J., & McArthur G. (2013). Validation of the Emotiv EPOC® ® EEG gaming system for measuring research quality auditory ERPs. PeerJ, 1, e38 DOI 10: 7717.

4. Behroozi M., Reza Daliri M., & Shekarchi B. (2015). EEG phase patterns reflect the representation of semantic categories of objects. Medical and Biological Engineering and Computing, DOI 10.1007/s11517–015–1391–7.

5. Boutani H., & Ohsuga M. (2013). Applicability of the Emotiv EEG Neuroheadset as a user—friendly input interface. Engineering in Medicine and Biology Society (EMBC), 2013 35th Annual International Conference of the IEEE, 1346–1349.

6. Croft R. J., & Barry R. J. (2000). Removal of ocular artifact from the EEG: a review. Clinical Neurophysiology, 30(1), 5–19.

7. Dixon W. J., & Tukey J. W. (1968). Approximate behavior of the distribution of Winsorized t (Trimming/Winsorization 2). Technometrics, 10(1), 83–98.

8. Hu L., Liang M., Mouraux A., Wise R. G., Hu Y., & Iannetti G. D. (2011). Taking into account latency, amplitude, and morphology: improved estimation of single–trial ERPs by wavelet filtering and multiple linear regression. Journal of Neurophysiology, 106(6), 3216–3229.

9. Jervis B.W., Ifeachor E.C., & Allen E.M. (1988). The removal of ocular artefacts from the electroencephalogram: a review. Medical and Biological Engineering and Computing, 26(1), 2–12.

10. Jin J., Allison B.Z, Sellers E.W., Brunner C., Horki P., Wang X., & Neuper C. (2011) Optimized stimulus presentation patterns for an event–related potential EEG–based brain–computer interface. Medical and Biological Engineering and Computing, 49(2), 181–191.

11. Joyce C. A., Gorodnitsky I. F., & Kutas M. (2004). Automatic removal of eye movement and blink artifacts from EEG data using blind component separation. Psychophysiology, 41(2), 313–325.

12. Linden D. E. (2005). The P300: where in the brain is it produced and what does it tell us? The Neuroscientist, 11(6), 563–576.

13. Liu Y, Jiang X., Cao T., Wan F., Mak P. U., Mak P. I., & Vai M. (2012, July). Implementation of SSVEP based BCI with Emotiv EPOC. Virtual Environments Human–Computer Interfaces and Measurement Systems (VECIMS), 2012 IEEE International Conference, 34–37.

14. Luck Steven J. (2005). An Introduction to the Event–Related Potential Technique. The MIT Press. <u>ISBN 0–262–12277–4</u>.

15. Makeig S., Bell A. J., Jung T. P., & Sejnowski T. J. (1996). Independent component analysis of electroencephalographic data. Advances in neural information processing systems, 8, 145–151.

16. Quiroga R. Q., & Garcia H. (2003). Single–trial event–related potentials with wavelet denoising. Clinical Neurophysiology, 114(2), 376–390.

17. Ramirez–Cortes J. M., Alarcon–Aquino V, Rosas–Cholula G., Gomez–Gil P., & Escamilla—Ambrosio J. (2010). P–300 rhythm detection using ANFIS algorithm and wavelet feature extraction in EEG signals. Proceedings of the World Congress on Engineering and Computer Science (Vol. 1, pp. 963–968). San Francisco: International Association of Engineers.

18. Sanei S., & Chambers J. A. (2013). EEG signal processing. John Wiley & Sons.

19. Sano A., & Bakardjian H. (2009). Movement–related cortical evoked potentials using four–limb imagery. International Journal of Neuroscience, 119(5), 639–663.

20. Vorobyov S., & Cichocki A. (2002). Blind noise reduction for multisensory signals using ICA and subspace filtering, with application to EEG analysis. Biological Cybernetics, 86(4), 293–303.

21. Wang Z., Maier A., Leopold D. A., Logothetis N. K., & Liang H. (2007). Single–trial evoked potential estimation using wavelets. Computers in Biology and Medicine, 37(4), 463–473.

22. Wastell D. G. (1977). Statistical detection of individual evoked responses: an evaluation of Woody’s adaptive filter. Electroencephalography and Clinical Neurophysiology, 42(6), 835–839.

